## supplementary information files for "Resolving subcellular pH with a quantitative fluorescent lifetime biosensor"

### Supplementary figures / videos:

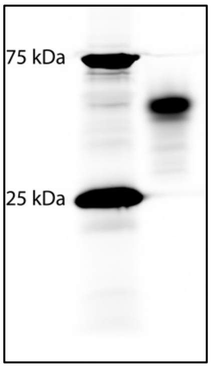


**Supplementary figure 1.** Fluorescent image of recombinant mApple protein (right) run on non-reducing SDS-PAGE to maintain mApple fluorescence. Molecular weight ladder shown on left.

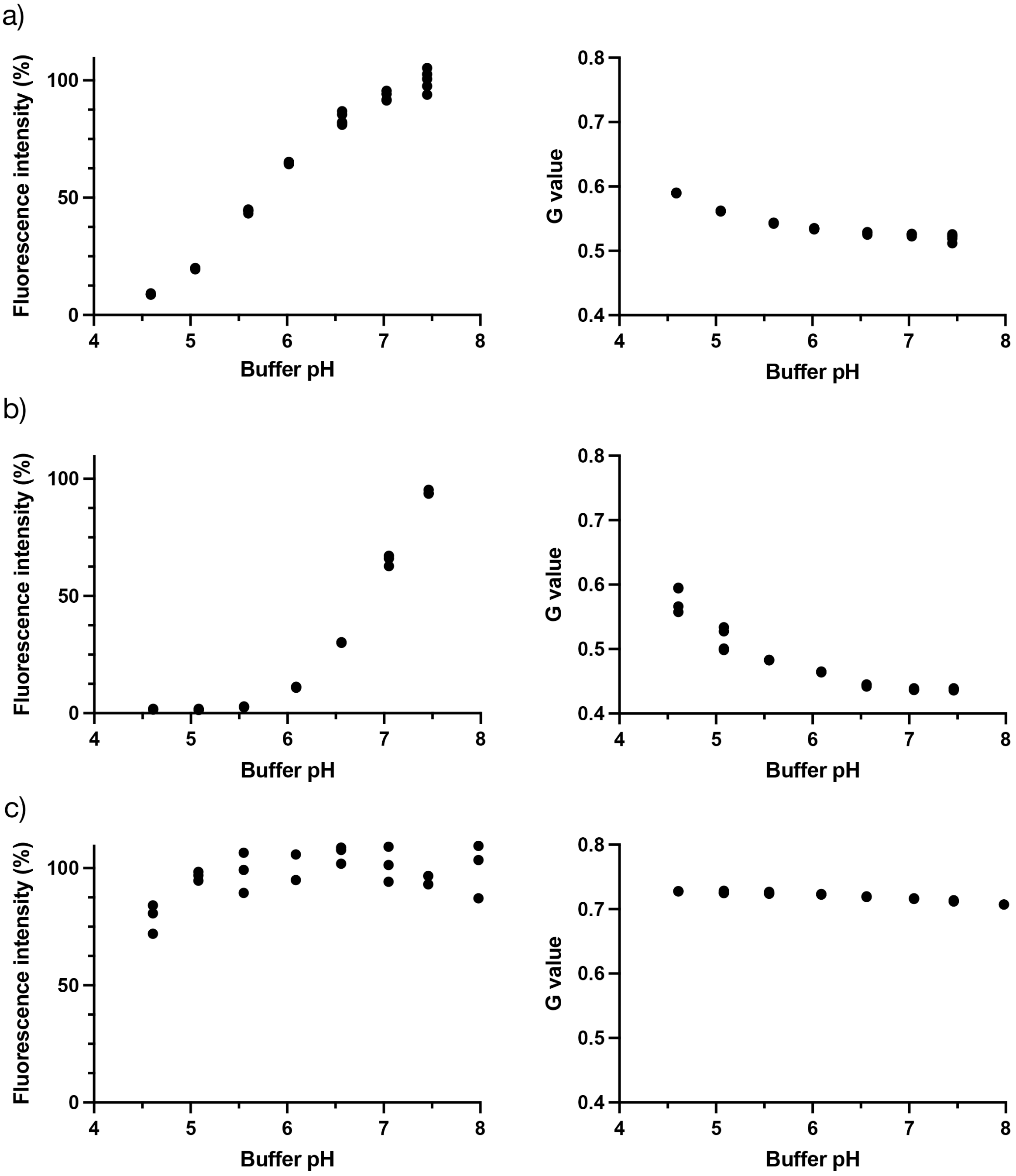


**Supplementary figure 2.** pH dependant fluorescence intensity (left) and lifetime (G value - right) of: **a**, muGFP; **b**, pHluorin; and **c**, mCherry. Points indicate mean, error bars of standard deviation are too small to be shown, n=3 independent replicates.

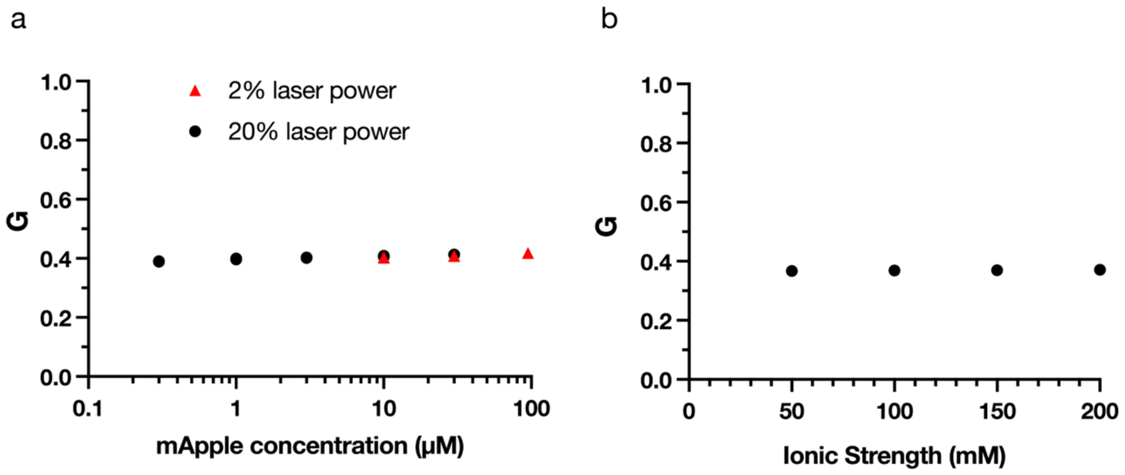


**Supplementary figure 3.** mApple concentration and ionic strength dependence. **a**, mApple concentration and laser power dependent effects on mApple lifetime at pH 7.3. There is minimal difference in the measured fluorescent lifetime for mApple protein at different concentrations and different laser powers. **b**, Influence of ionic strength on mApple G value. No variability is observed within the indicated ionic strengths. Points indicate mean, error bars of standard deviation are too small to be shown, n=3 independent replicates.

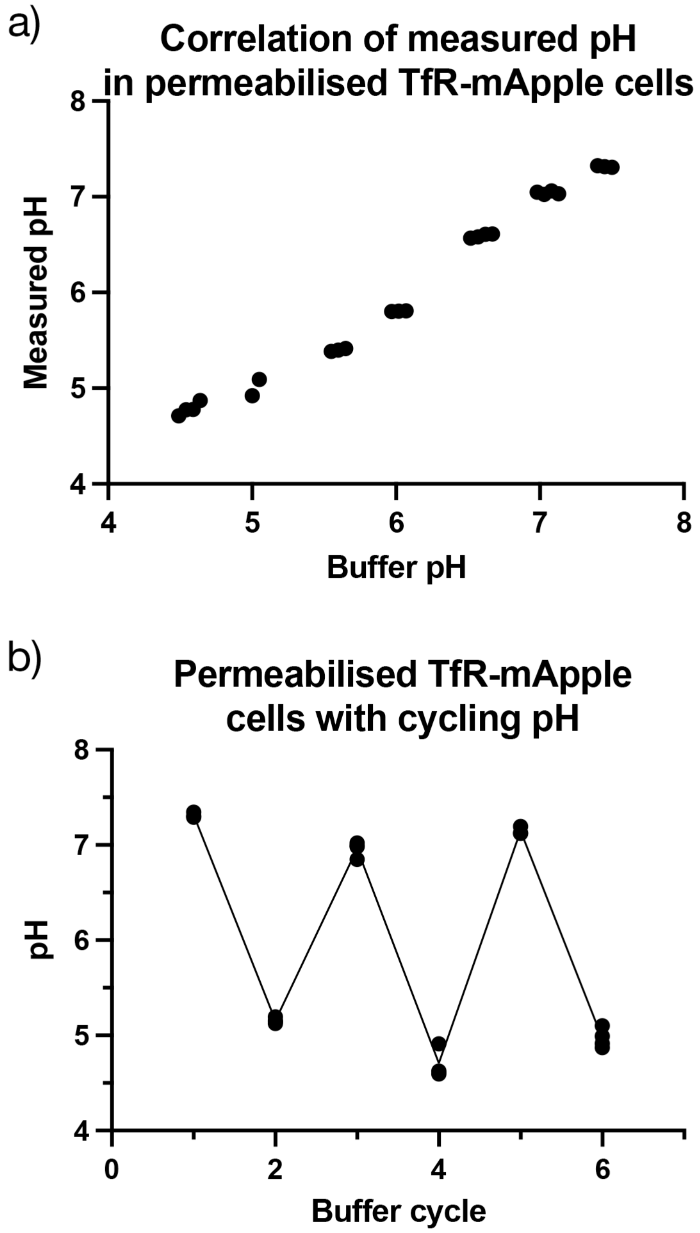


**Supplementary figure 4.** NIH-3T3 cells expressing TfR-mApple permeabilised with 0.02% digitonin and incubated with different pH buffers. **a**, Correlation of measured pH (using free mApple calibration curve – Figure 1g) with buffer pH. The slight deviation from linear behaviour is likely due to incomplete permeabilization of the cells and difficulty equilibrating the pH throughout the entire cell. **b**, Cycling of pH between 7.4 and 4.6. Points indicate mean of entire image, n=3 independent replicates.

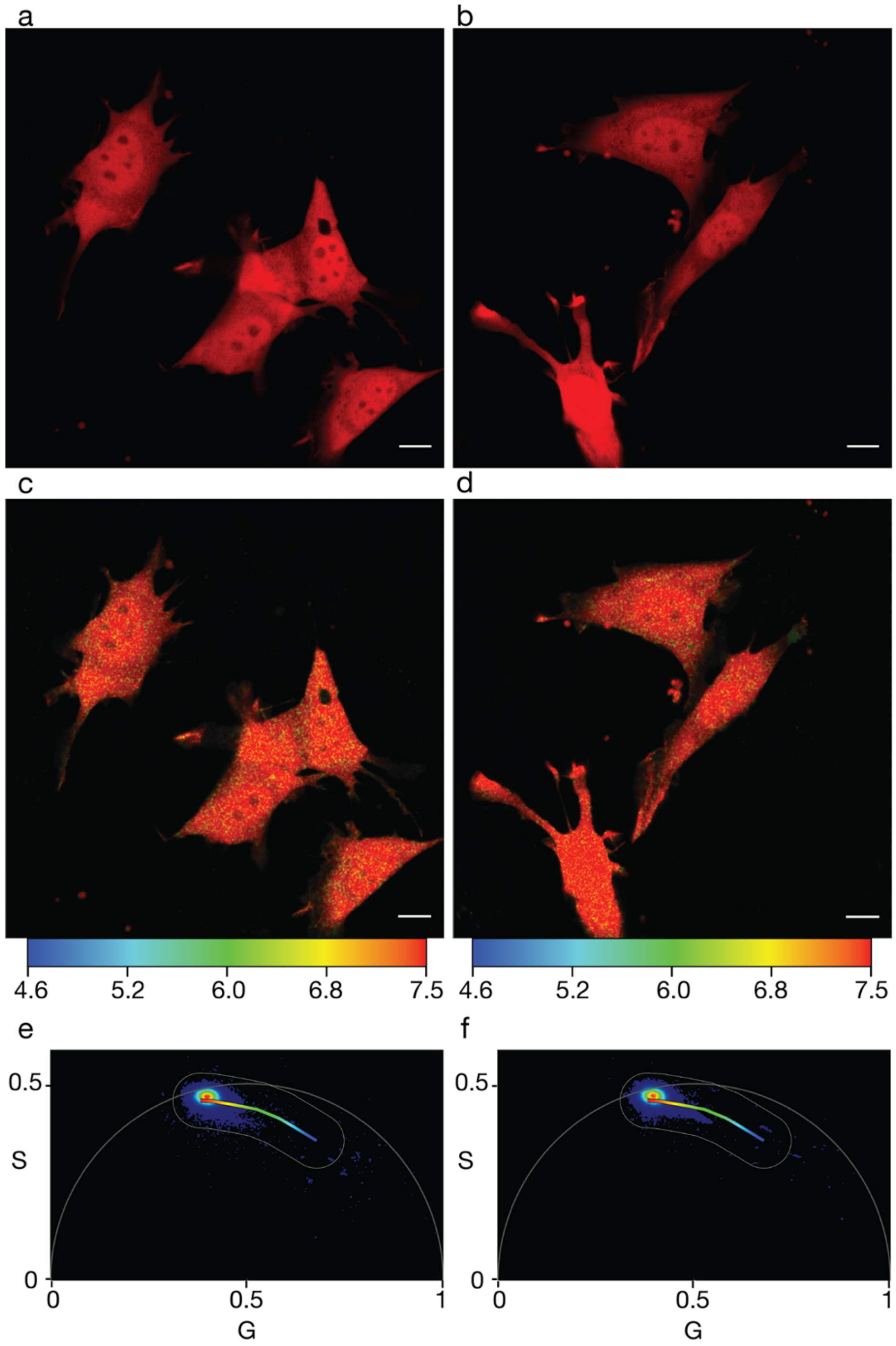


**Supplementary figure 5**. Additional images of NIH-3T3 cells expressing untagged mApple**. a,b**, Confocal images pseudo-coloured red. **c,d**, Corresponding fast FLIM images of a,b pseudo-coloured according to their pH. pH colour scale indicated underneath images. **e,f**, Corresponding phasor plots of c,d respectively, with an overlayed phasor mask. Phasor plot colour is indicative of the frequency of photons at that phasor position (red = high, blue = low). Scale bar = 10 μm.

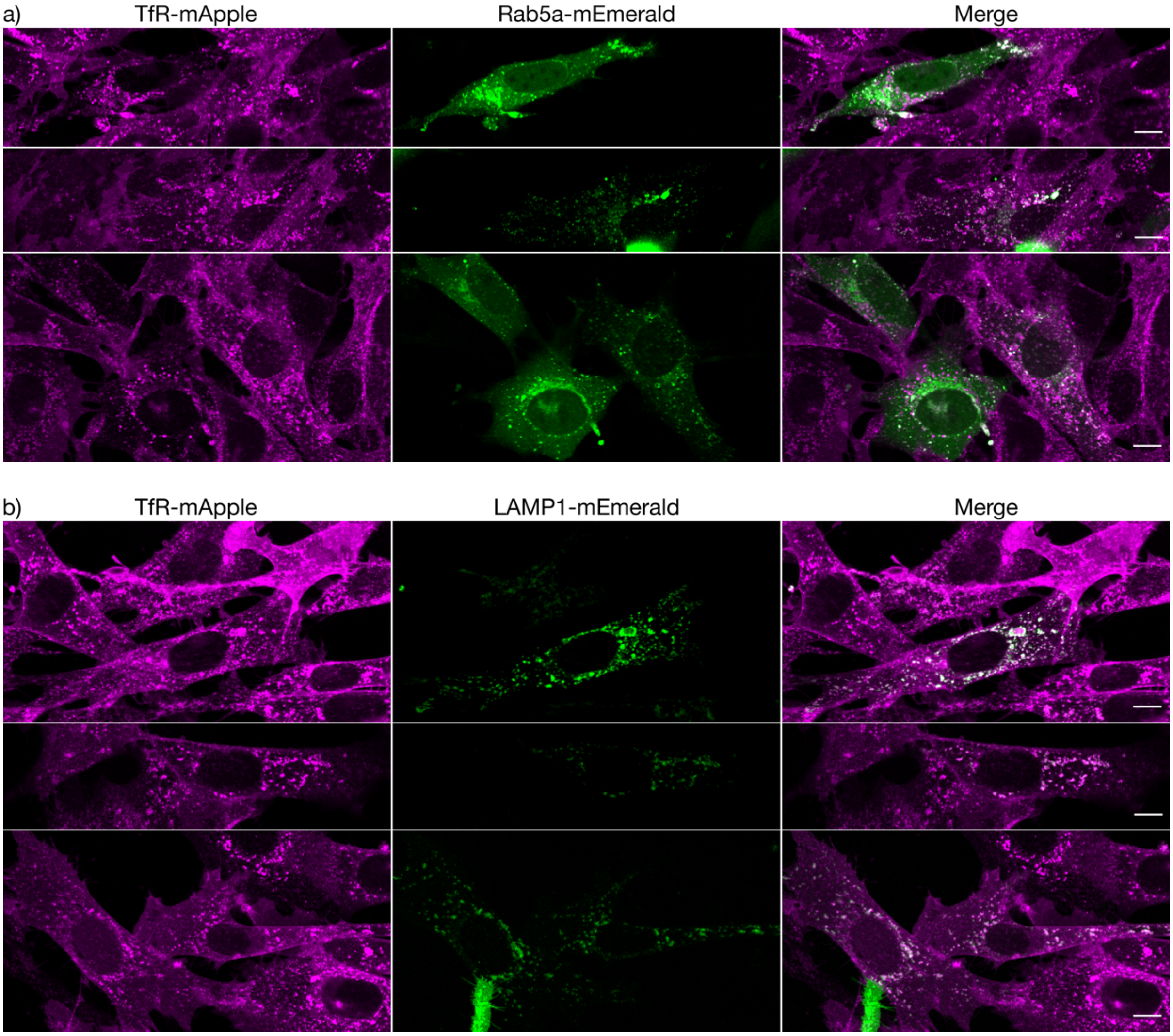
a)
TfR-mApple
Rab5a-mEmerald
Merge
b)
TfR-mApple
LAMP1-mEmerald
Merge

c)

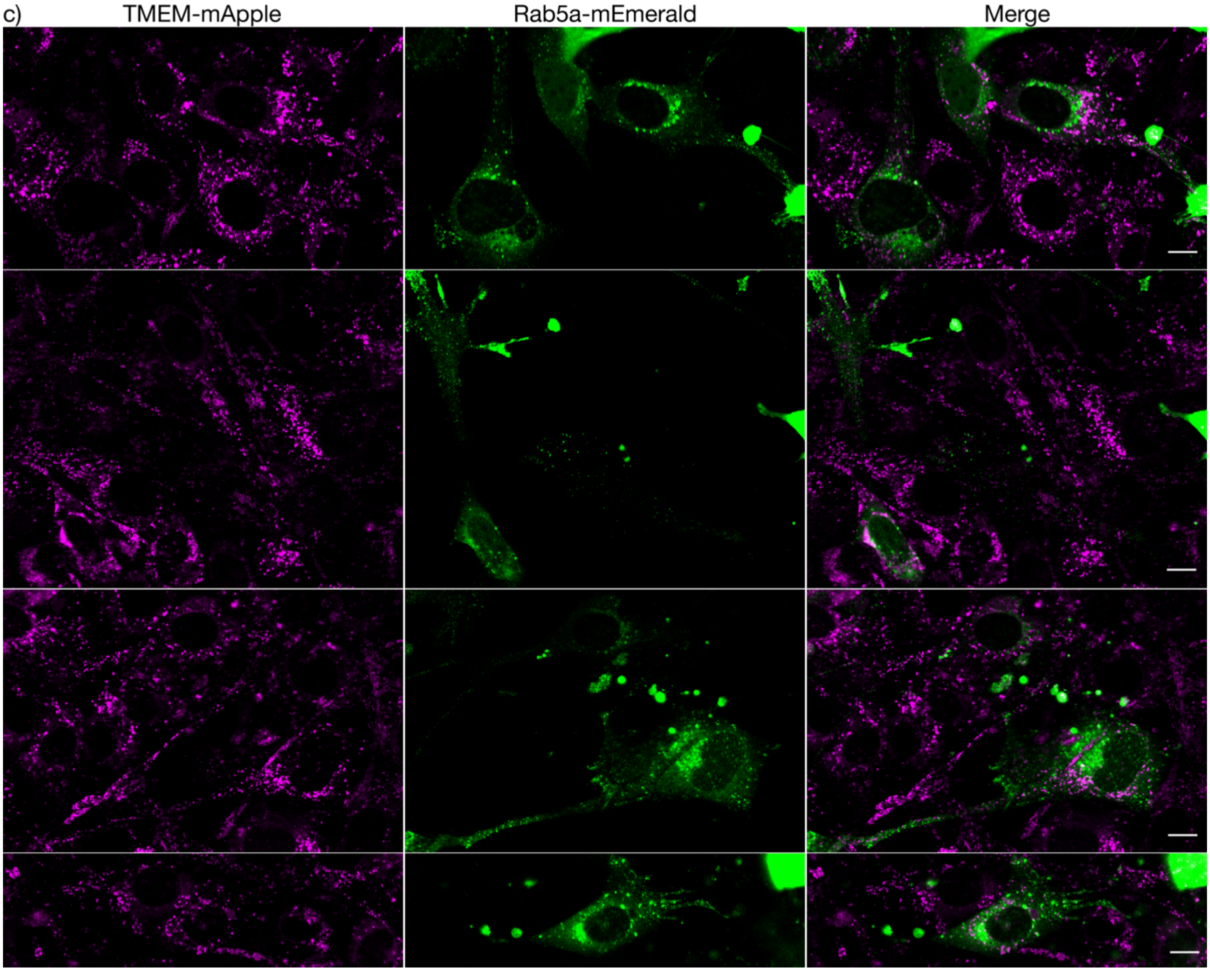

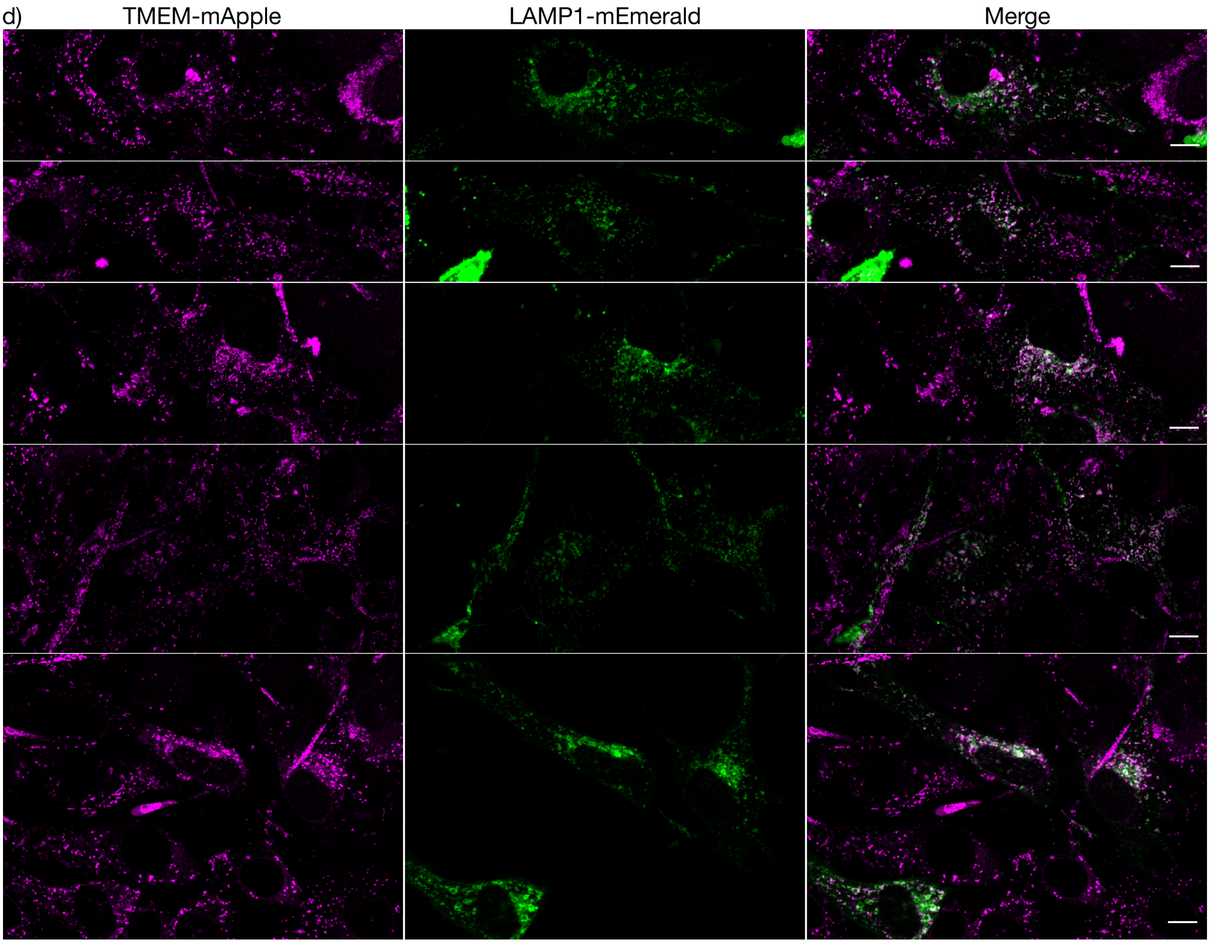


**Supplementary figure 6.** Colocalisation images of mApple fusions (pseudo-coloured magenta, column 1) with endosomal stage markers (pseudo-coloured green, column 2) and merge (column 3). Endosomal stage markers were transiently transfected into cell lines stably expressing mApple fusions. **a**, TfR-mApple with Rab5a-mEmerald. **b**, TfR-mApple with LAMP1-mEmerald. **c**, TMEM-mApple with Rab5a-mEmerald. **d**, TMEM-mApple with LAMP1-mEmerald. Cropped images are shown to represent Rab5a / LAMP1 transiently transfected cells in the field of view. Note, cells originate from stable lines selected for mApple expression, so all cells are mApple positive, however the transient transfection with mEmerald means that not all cells are positive for mEmerald. Scale bar = 10 μm.

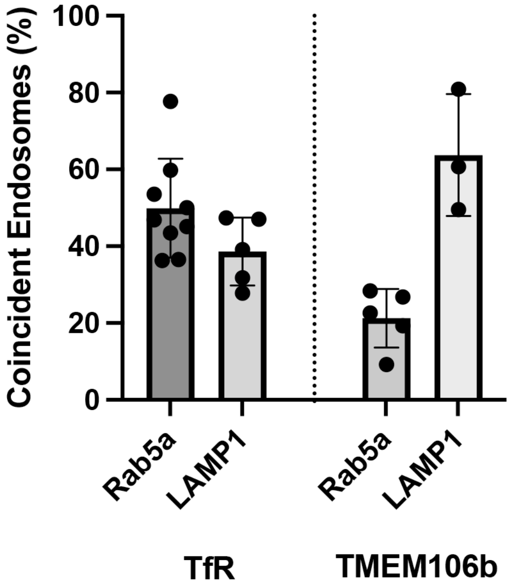


**Supplementary figure 7.** Coincidence analysis of mApple fusions shown in supplementary figure 6. Double positive cells were hand segmented and the StarDist algorithm was used to identify the mApple positive and mEmerald (indicating Rab5a or LAMP1) endosomes. The total number of double positive mApple/mEmerald endosomes was calculated and expressed as a percent of the total number of mApple positive endosomes. n>3 independent replicates.

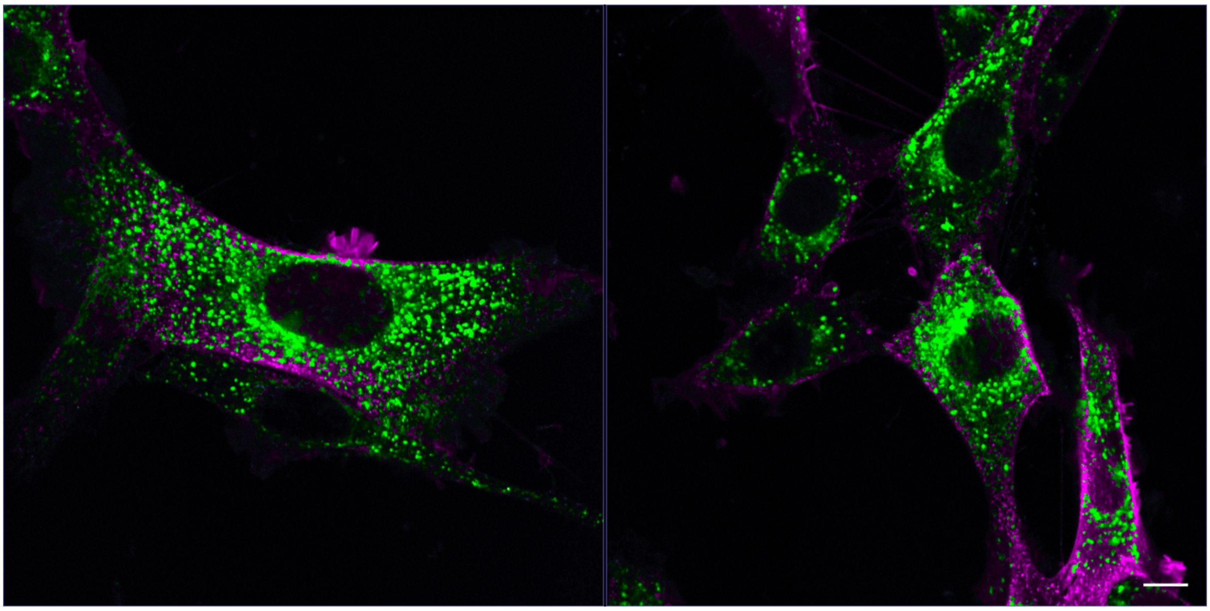

**Supplementary figure 9.** NIH-3T3 cells expressing mApple-TfR fusion protein. False coloured images with areas >pH 6.5 coloured magenta, and areas <pH 6.5 coloured green. High pH (red) areas are clearly visible on the surface of the cell, and lower pH (green) areas are visible inside the cell. Large patches of red are indicative of the imaging plane coinciding with the top of the cell. Scale bar = 10 μm.

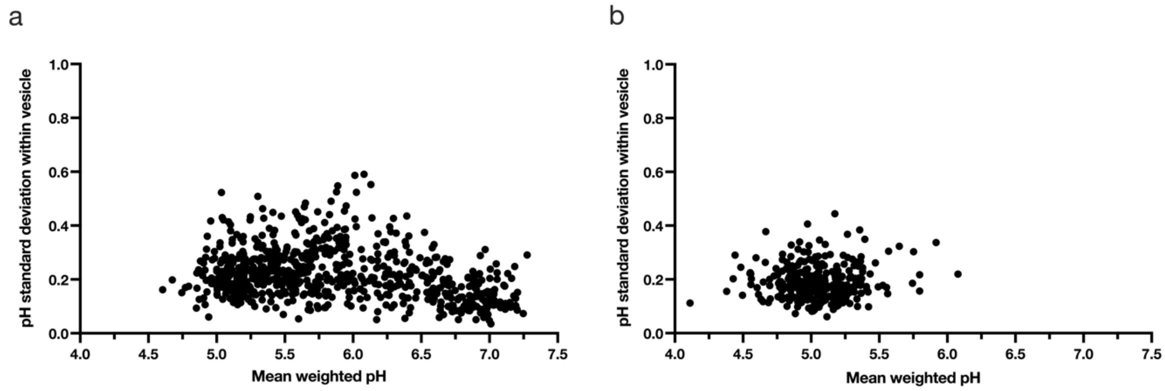


**Supplementary figure 10.** Comparison of pH variability within vesicles. **a,b**, Plot of pH standard deviation of vesicles detected in Fig. 3a (TfR-mApple) and Fig. 3c (TMEM-mApple) respectively.

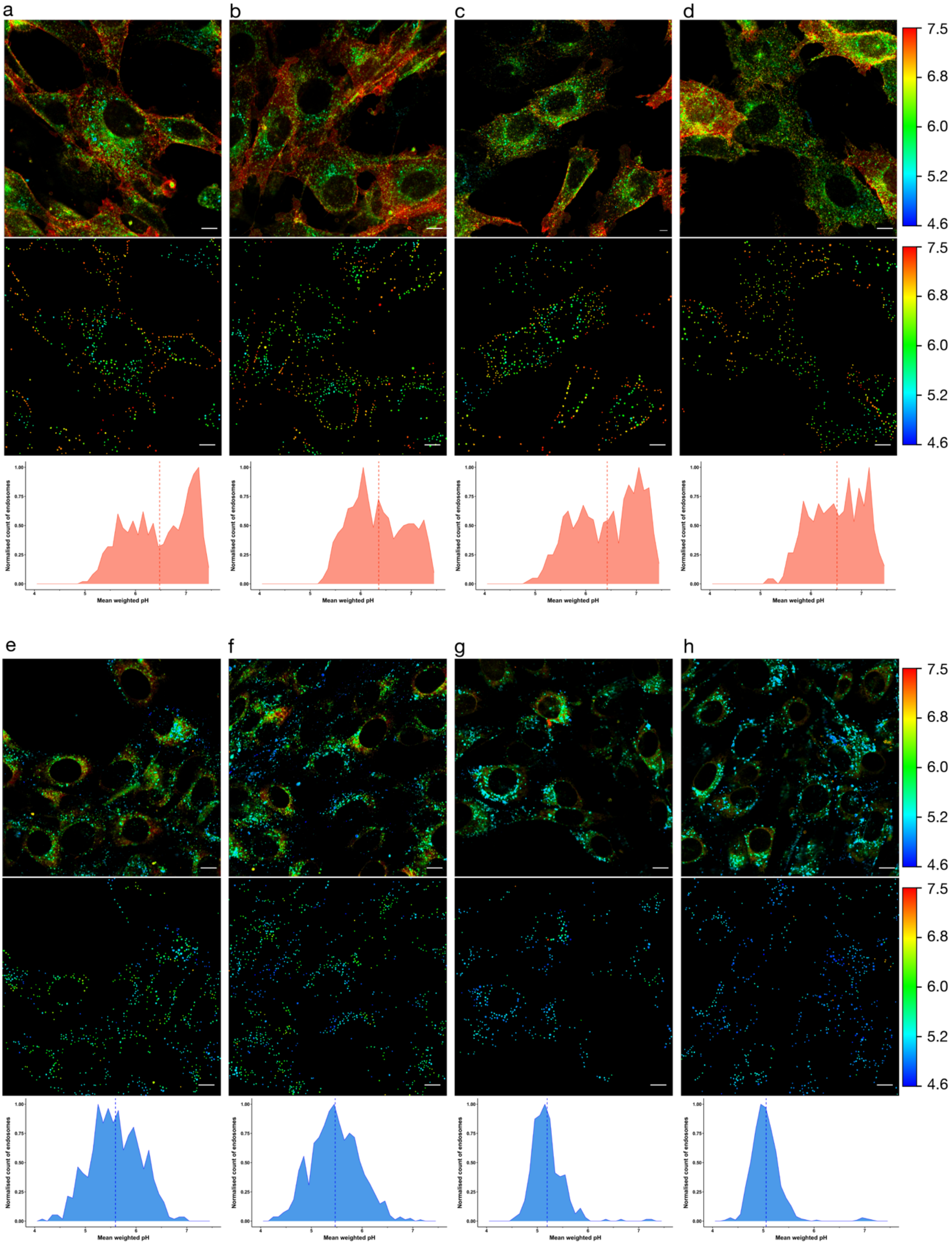

a
b
c
d
7.5
6.8
6.0
5.2
4.6
7.5
6.8
6.0
5.2
4.6
Normalised count of endosomes
Mean weighted pH
e
f
g
h
7.5
6.8
6.0
5.2
4.6
7.5
6.8
6.0
5.2
4.6
Normalised count of endosomes
Mean weighted pH

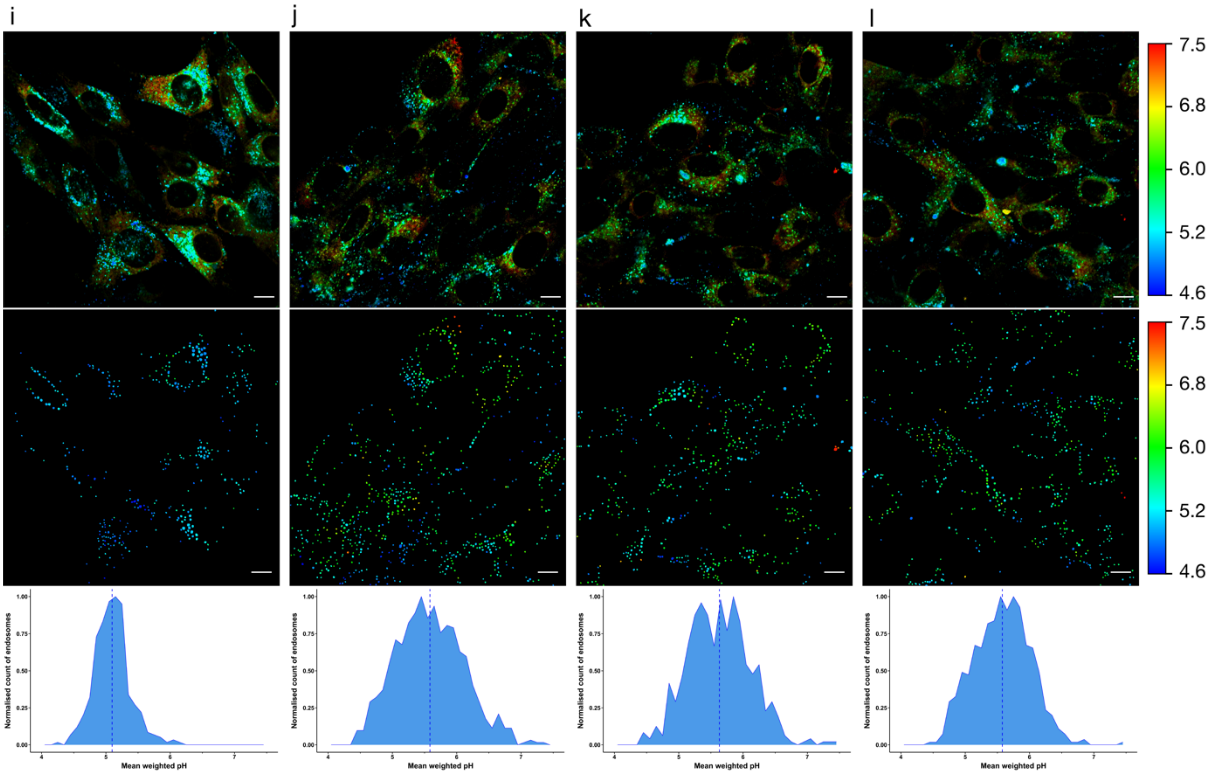


**Supplementary figure 11**. Visualising the pH of intracellular vesicles in NIH-3T3 cells using fast FLIM and automated endo/lysosomal compartment detection and analysis. **a-d**, TfR-mApple. **e-l**, TMEM-mApple. Top panel shows standard fluorescent intensity images pseudo coloured by a phasor mask (Fig. 1g) which corresponds to colour scale (right). Middle panel indicates automatically detected endo/lysosomes pseudo-coloured according to the mean weighted pH of the vesicle, corresponds to colour scale (right). Bottom panel displays each of the detected vesicles (middle panel) on a frequency histogram showing the distribution of endo/lysosomal mean weighted pH, population mean shown as a dotted line. Scale bar = 10 μm.

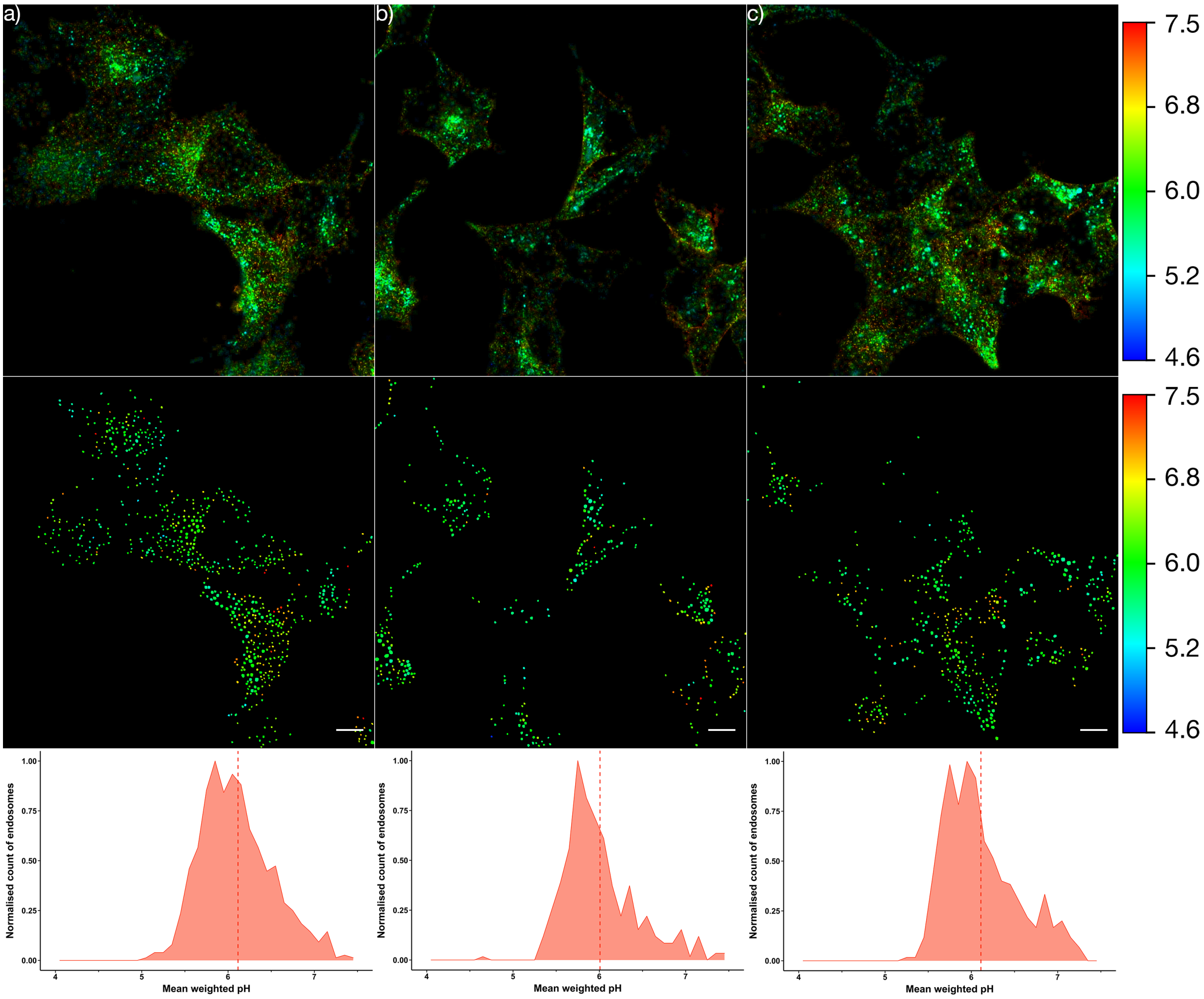


**Supplementary figure 12**. Visualising the pH of intracellular vesicles in HEK293 cells using fast FLIM and automated endo/lysosomal compartment detection and analysis. **a-c**, TfR-mApple. Top panel shows standard fluorescent intensity images pseudo coloured by a phasor mask (Fig. 1g) which corresponds to colour scale (right). Middle panel indicates automatically detected endo/lysosomes pseudo-coloured according to the mean weighted pH of the vesicle, corresponds to colour scale (right). Bottom panel displays each of the detected vesicles (middle panel) on a frequency histogram showing the distribution of endo/lysosomal mean weighted pH, population mean shown as a dotted line. Scale bar = 10 μm.

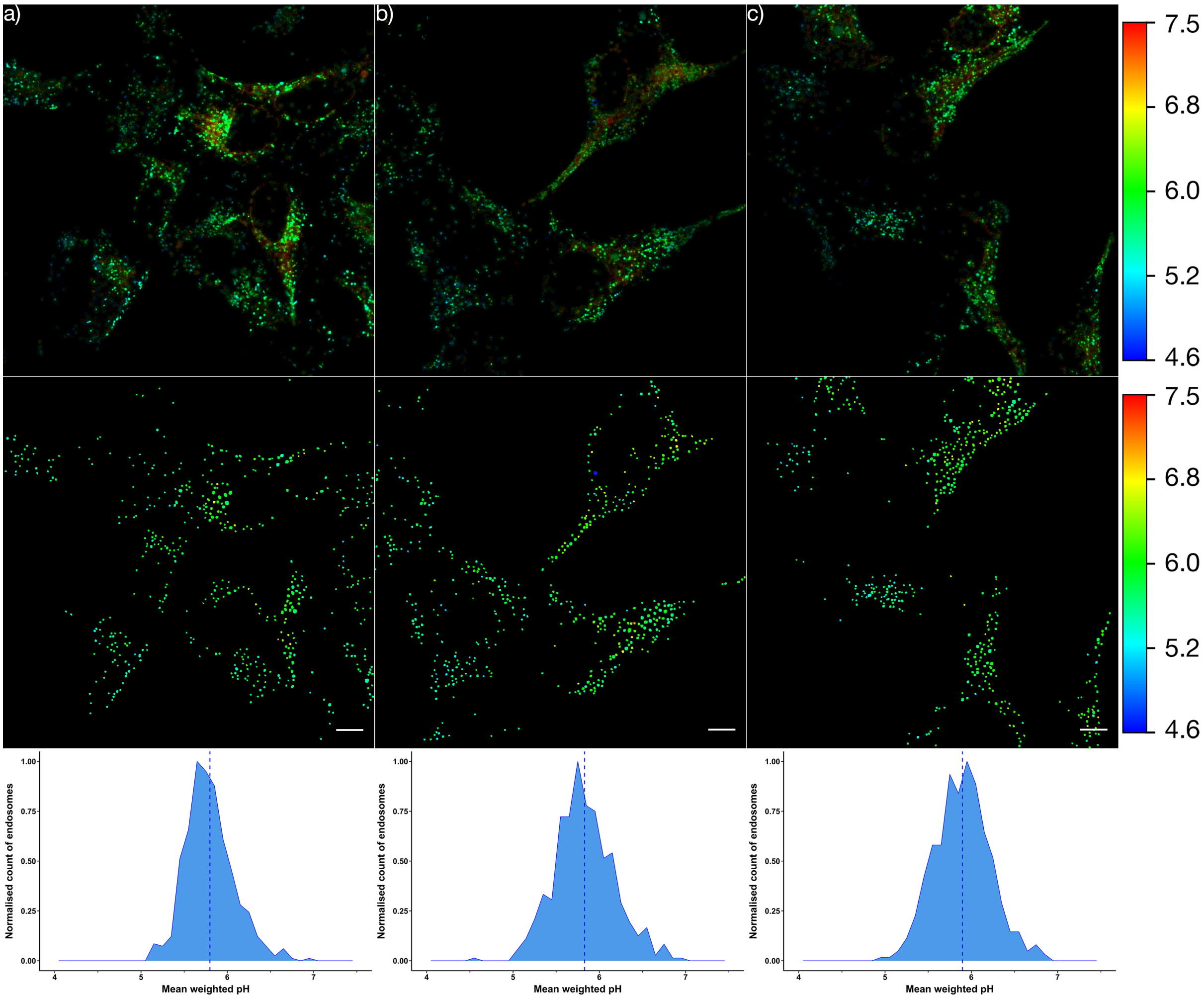


**Supplementary figure 13**. Visualising the pH of intracellular vesicles in HEK293 cells using fast FLIM and automated endo/lysosomal compartment detection and analysis. **a-c**, TMEM106b-mApple. Top panel shows standard fluorescent intensity images pseudo coloured by a phasor mask (Fig. 1g) which corresponds to colour scale (right). Middle panel indicates automatically detected endo/lysosomes pseudo-coloured according to the mean weighted pH of the vesicle, corresponds to colour scale (right). Bottom panel displays each of the detected vesicles (middle panel) on a frequency histogram showing the distribution of endo/lysosomal mean weighted pH, population mean shown as a dotted line. Scale bar = 10 µm.

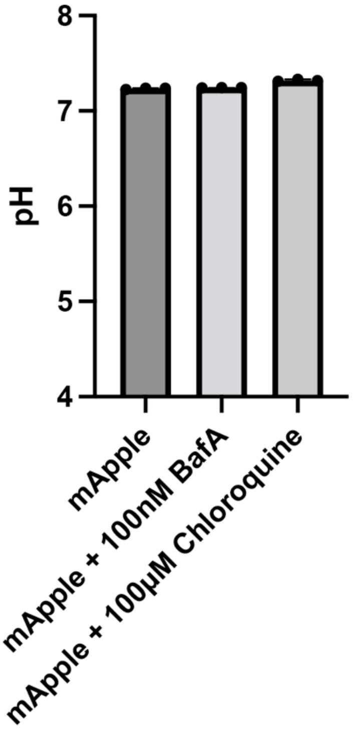


**Supplementary figure 14.** Effect of BafA and Chloroquine on the lifetime of mApple. mApple protein was incubated with 100nM BafA or 100µM of Chloroquine for 45 min prior to imaging. The addition of either chemical did not affect the pH measurement inferred from mApple fluorescent lifetime.

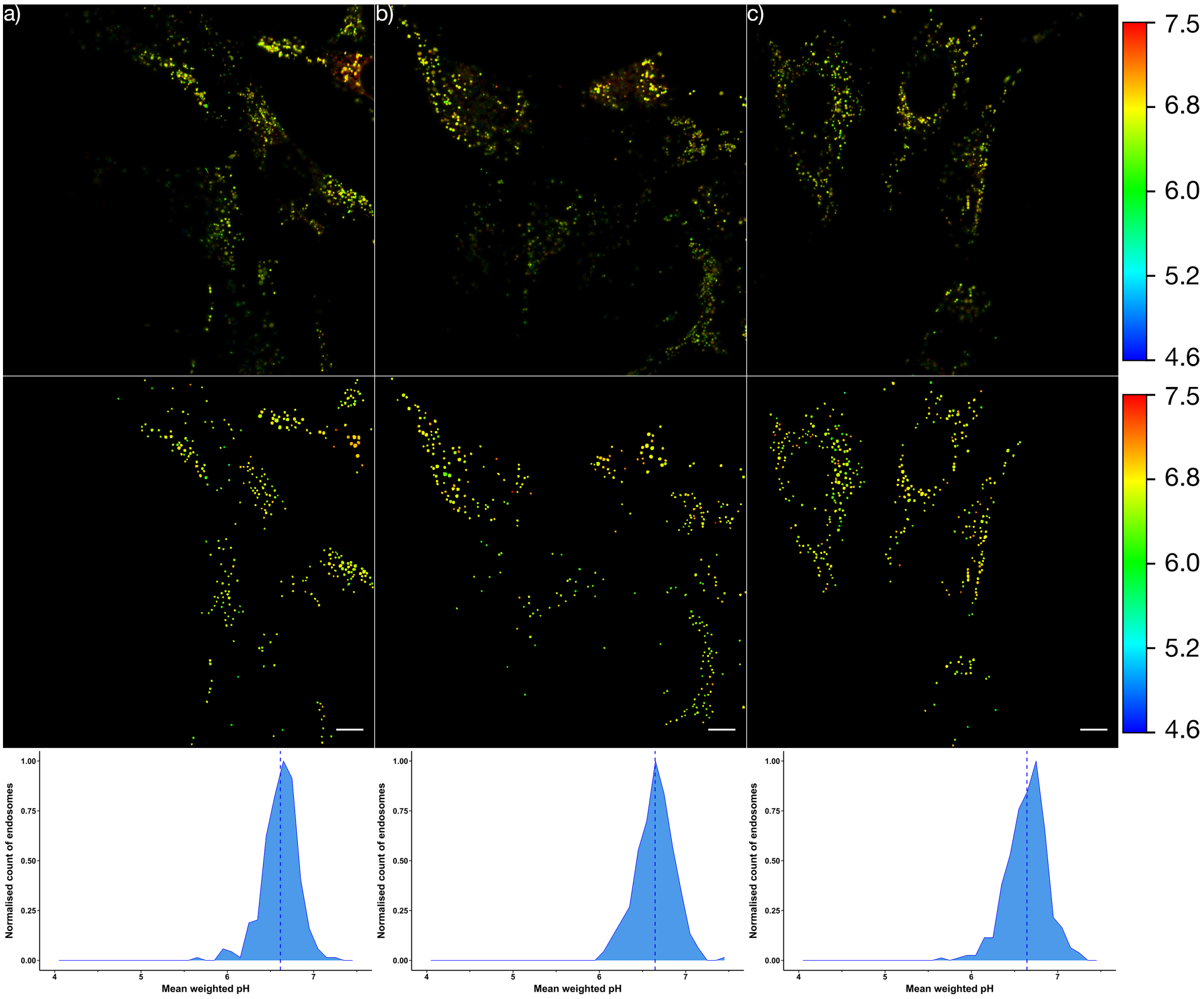


**Supplementary figure 15.** Visualising the pH of intracellular vesicles in NIH-3T3 cells expressing TMEM106b-mApple treated with 100µM chloroquine for 45 min. Top panel shows standard fluorescent intensity images pseudo coloured by a phasor mask (Fig. 1g) which corresponds to colour scale (right). Middle panel indicates automatically detected endo/lysosomes pseudo-coloured according to the mean weighted pH of the vesicle, corresponds to colour scale (right). Bottom panel displays each of the detected vesicles (middle panel) on a frequency histogram showing the distribution of endo/lysosomal mean weighted pH, population mean shown as a dotted line. Scale bar = 10 µm.

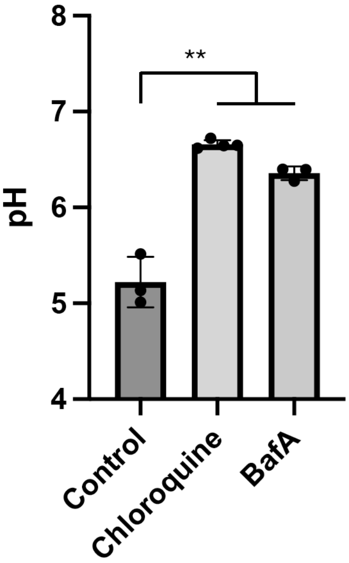


**Supplementary figure 16.** Comparison of the average endosomal pH of NIH-3T3 cells expressing TMEM106b-mApple and treated with either 100μM of chloroquine or 100nM of BafA for 45 min. Treatment with both chloroquine or BafA resulted in a significant increase in endosomal pH (** denotes *p* value < 0.01).

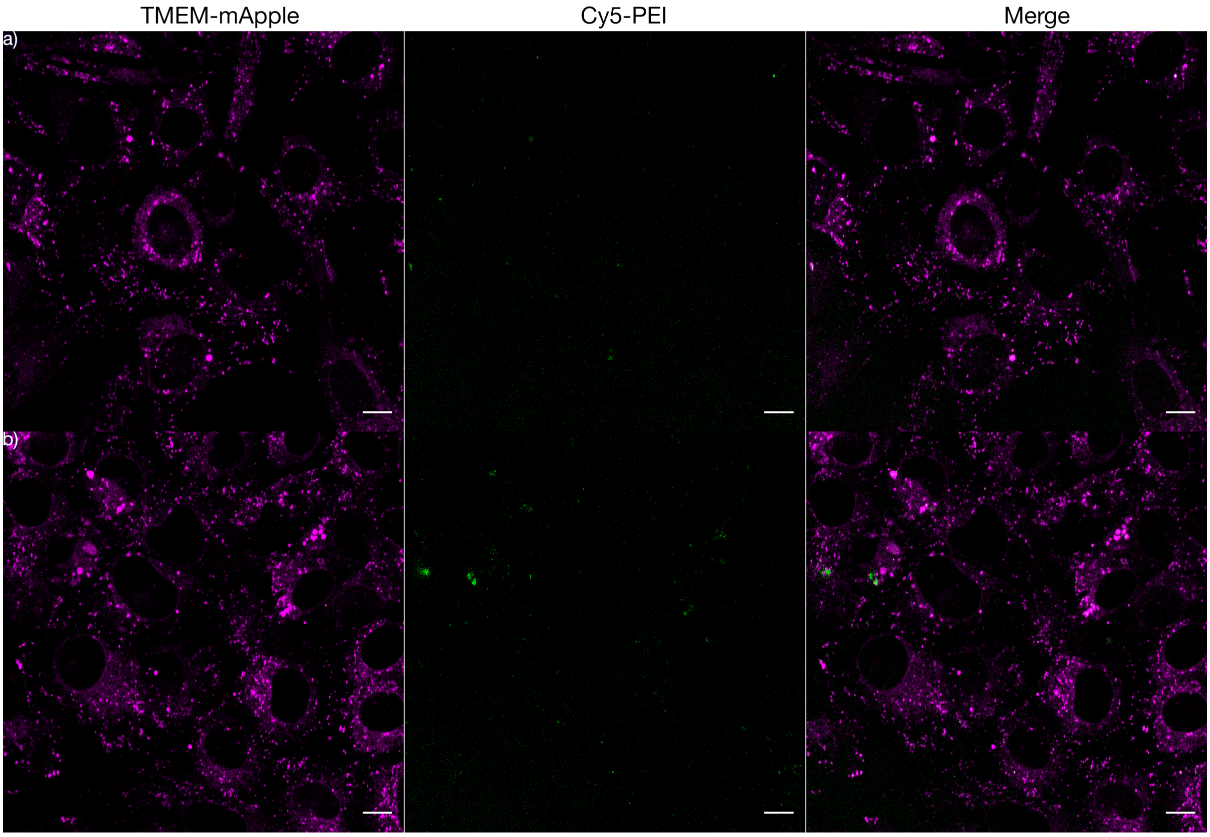

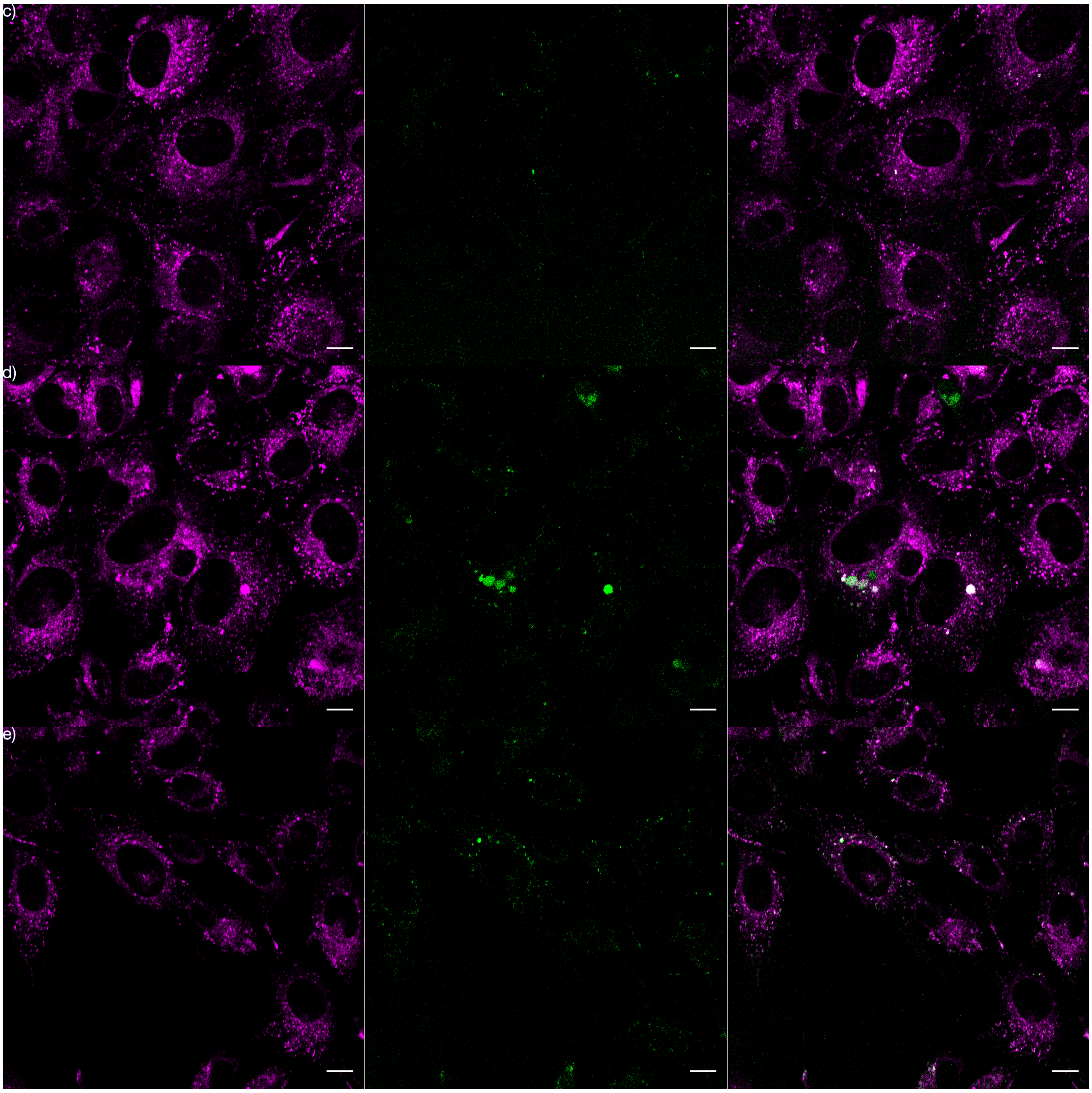
c)
d)
e)

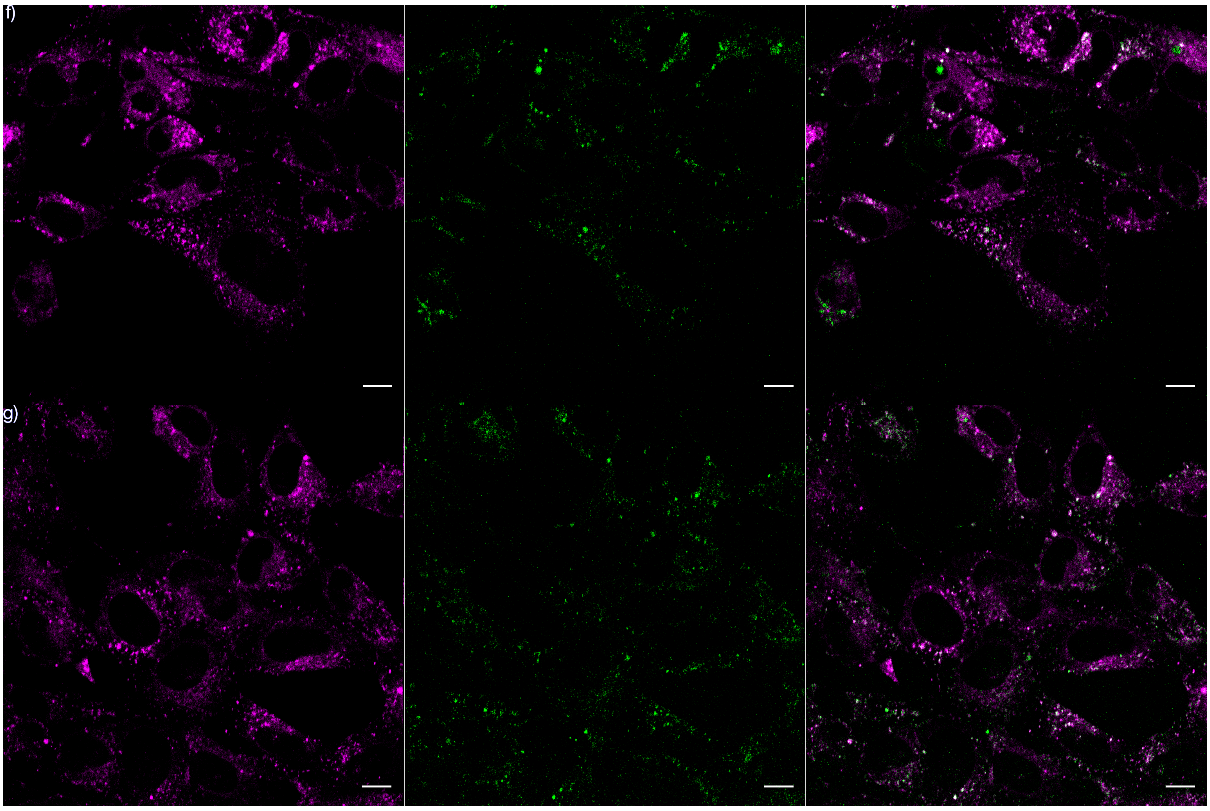


**Supplementary figure 17.** Time course of NIH-3T3 cells treated with PEI labelled with Cy5. TMEM106b-mApple pseudo coloured magenta (panel 1), Cy5-PEI pseudo coloured green (panel 2) and merge (panel 3). Colocalisation is observed after 60 minutes. **a**, 0 min, **b**, 30 min, **c**, 60 min, **d**, 120 min, **e**, 240 min, **f**, 300 min, **g**, 360 min. n=1.

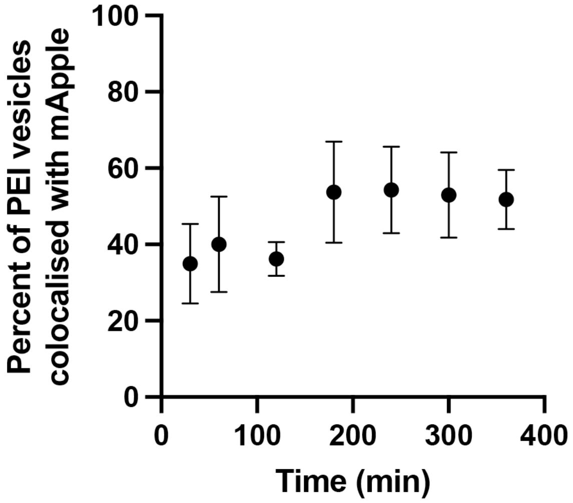


**Supplementary figure 18.** Coincidence analysis of TMEM106b-mApple expressed in NIH-3T3 cells with Cy5 labelled PEI. The StarDist algorithm was used to identify the mApple positive and Cy5 positive endosomes. The total number of double positive mApple/Cy5 endosomes was calculated and expressed as a percent of the total number of Cy5 positive endosomes. n>3 independent replicates.

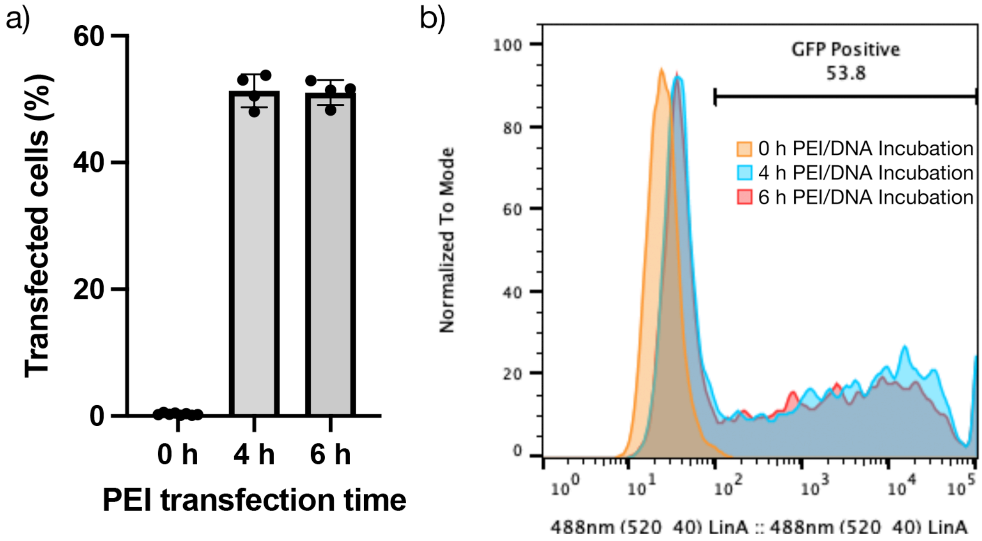


**Supplementary figure 19.** Transfection efficiency of NIH-3T3 cells expressing mApple-TMEM106b incubated with Cy5 labelled PEI/pDNA complexes. Polyplexes were assembled with Cy5 labelled and transfection grade PEI (ratio 1:5, 80µg/mL final) in addition to pDNA encoding EGFP (2µg/mL final), which were incubated with cells for 0, 4, and 6 hours before the cells were washed. EGFP fluorescence was assessed by flow cytometry 24 hours post PEI addition.

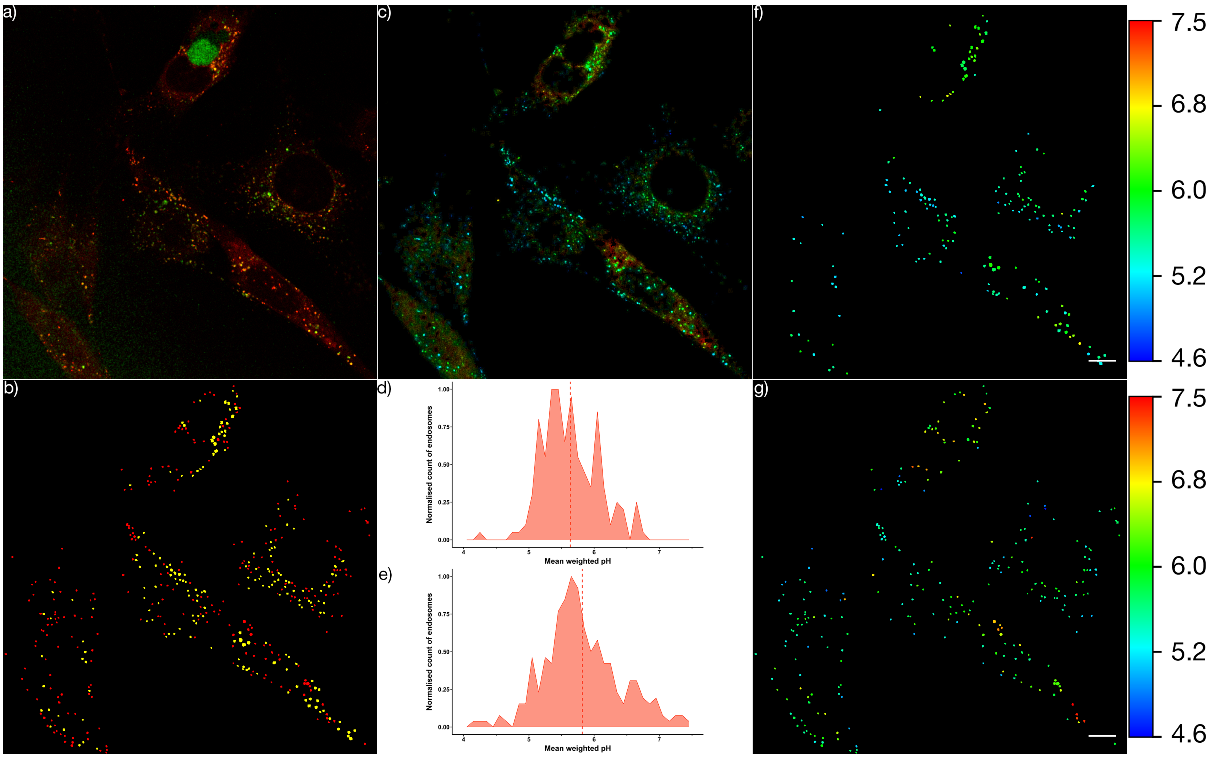


**Supplementary figure 20.** Visualising the pH of intracellular vesicles in NIH-3T3 cells expressing TMEM106b-mApple treated with Cy5 labelled PEI/DNA complexes. **a**, Overlay of mApple (red) and Cy5 (green) signal. **b**, vesicles identified by the StarDist algorithm: red mApple only, yellow double positive for mApple and Cy5. Cy5 only vesicles not shown. **c**, mApple fluorescent intensity image pseudo coloured by a phasor mask (Fig. 1g) which corresponds to colour scale (right). **d**, frequency histogram showing the distribution of endo/lysosomal mean weighted pH of vesicles that contain PEI. **e**, frequency histogram showing the distribution of endo/lysosomal mean weighted pH of vesicles that do not contain PEI. **f,g**, automatically detected endo/lysosomes pseudo-coloured according to the mean weighted pH of the vesicle, corresponds to colour scale (right). **f**, endosomes with PEI. **g**, endosomes without PEI. Scale bar = 10 μm.

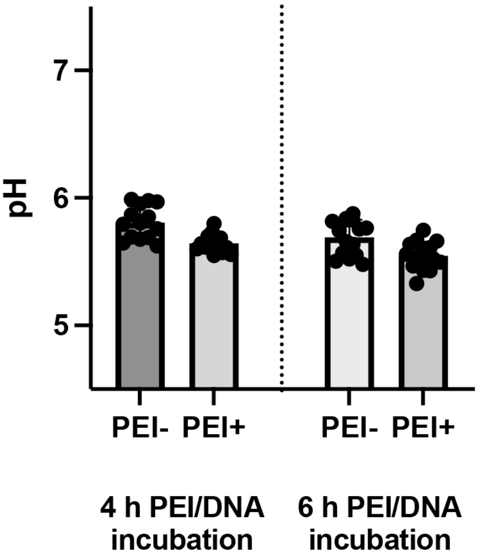


**Supplementary figure 21.** Average pH of endosomes in NIH-3T3 cells expressing mApple-TMEM106b and treated with Cy5 labelled PEI/DNA complexes. mApple-TMEM106b endosomes with or without PEI were detected as outlined in S20. n>10 images.

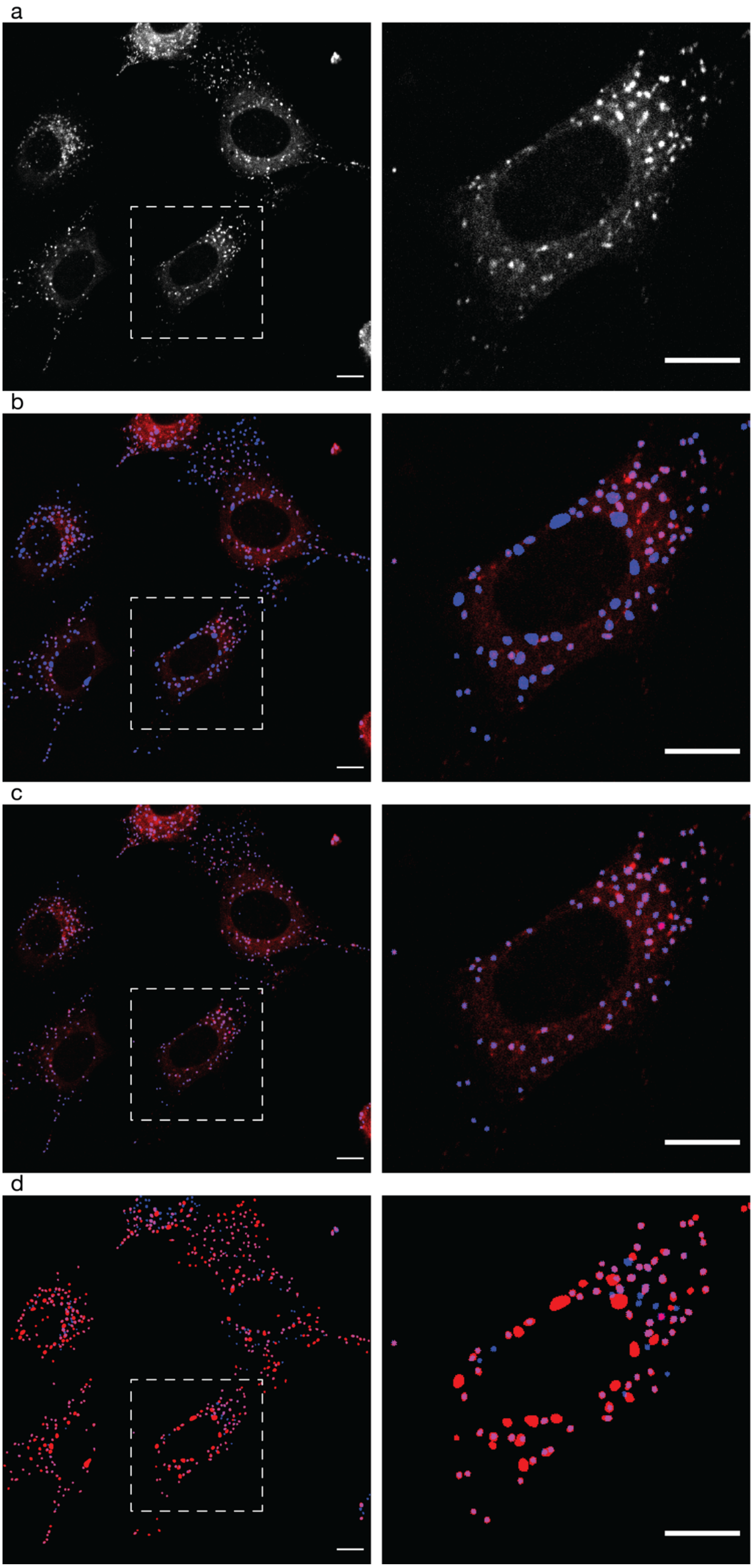

a
b
c
d

**Supplementary figure 22.** New model detects the majority of intracellular vesicles whilst eliminating the occurrence of false positives. **a**, TMEM-mApple fluorescent image used for algorithm comparison. **b**, Applying the versatile (fluorescent nuclei) stardist algorithm, TMEM-mApple fluorescence pseudo-coloured red, algorithm output (detected vesicles) pseudo-coloured blue. **c**, Applying the retrained algorithm, TMEM-mApple fluorescence pseudo-coloured red, algorithm output (detected vesicles) pseudo-coloured blue. **d**, Comparison of algorithm outputs, versatile (fluorescent nuclei) (b) pseudo-coloured red, retrained algorithm (c) pseudo-coloured blue. Inset shown on right, scale bar = 10µm in all images. Algorithm settings were as per methods section for both models.

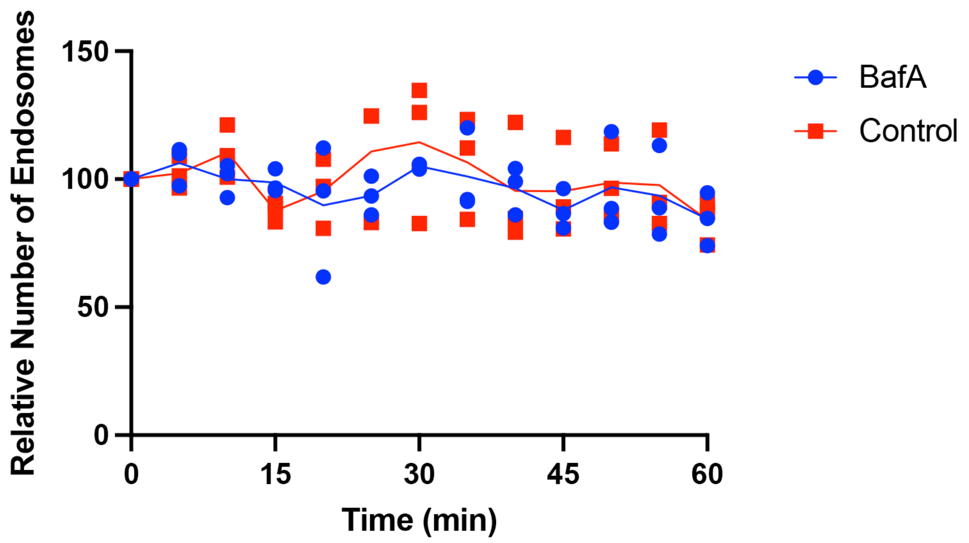


**Supplementary figure 23.** Relative number of endosomes detected in NIH-3T3 cells expressing mApple-TMEM106b and treated with 100nM BafA over 60 min. The same field of view was imaged over 60 minutes (see Figure 4) and the number of vesicles detected in each field of view was normalised to 100 for the 0 time point. n=3 independent replicates.

**Supplementary movie 1.** Timelapse of TfR-mApple expressing cell over 75 seconds at 1.4 second intervals. mApple intensity shown, phasor mask applied for pseudo-colouring pH according to the G-value calibration (Fig. 1g). Scale bar = 10µm.

**Supplementary movie 2.** Time course of TMEM106b-mApple expressing cells either bafilomycin A1 (100nM) treated (**a**) or untreated (**b**). Left panel sows mApple intensity with a phasor mask applied for pseudo-colouring pH according to the G-value calibration (Fig. 1g). Middle panel identifies automated detection of intracellular vesicles, pseudo coloured according to the vesicle mean weighted pH (shown inset). Scale bars = 10µm. Right panel shows the corresponding phasor plot with overlayed phasor mask (Fig. 1g). Time indicated top right.
